# Pherotype polymorphism in *Streptococcus pneumoniae* and its effects on population structure and recombination

**DOI:** 10.1101/070011

**Authors:** Eric L. Miller, Benjamin A. Evans, Omar E. Cornejo, Ian S. Roberts, Daniel E. Rozen

**Affiliations:** Faculty of Biology, Medicine, and Health, University of Manchester, Manchester, United Kingdom; Institute of Biology, Leiden University, Leiden, Netherlands; Norwich Medical School, University of East Anglia, Norwich, United Kingdom; School of Biological Sciences, Washington State University, Pullman, USA

**Author notes:** Corresponding authors: Ian S. Roberts, Oxford Road, Manchester M13 9PL, England, The United Kingdom, Daniel E. Rozen, Sylviusweg 72, 2333 BE Leiden, The Netherlands.

**Keywords:** competence, pneumococcus, quorum sensing, balanced polymorphism, horizontal gene transfer

## Abstract

Natural transformation in the Gram-positive pathogen *Streptococcus pneumoniae* occurs when cells become “competent”, a state that is induced in response to high extracellular concentrations of a secreted peptide signal called CSP (Competence Stimulating Peptide) encoded by the *comC* locus. Two main CSP signal types (pherotypes) are known to dominate the pherotype diversity across strains. Using thousands of fully sequenced pneumococcal genomes, we confirm that pneumococcal populations are highly genetically structured and that there is significant variation among diverged populations in pherotype frequencies; most carry only a single pherotype. Moreover, we find that the relative frequencies of the two dominant pherotypes significantly vary within a small range across geographical sites. It has been variously proposed that pherotypes either promote genetic exchange among cells expressing the same pherotype, or conversely that they promote recombination between strains bearing different pherotypes. We distinguish these hypotheses using a bioinformatics approach by estimating recombination frequencies within and between pherotypes across 4,089 full genomes. Despite underlying population structure, we observe extensive recombination between populations; additionally, we found significantly higher rates of genetic exchange between strains expressing different pherotypes than among isolates carrying the same pherotype. Our results indicate that pherotypes do not restrict, and marginally facilitate, recombination between strains. Furthermore, our results suggest that the CSP balanced polymorphism does not causally underlie population differentiation. Therefore, when strains carrying different pherotypes encounter one another during co-colonization, genetic exchange can freely occur.

## Introduction

The Gram-positive pathogen *Streptococcus pneumoniae* is responsible for up to 1 million deaths annually (O’Brien et al., 2009). *S. pneumoniae* is naturally transformable, and this ability to take up and recombine extracellular DNA across a broad size range, from 15 bp to 19 kb (Mostowy et al., 2014), is associated with the acquisition of antibiotic resistance genes and with capsular switching (Croucher et al., 2014a, 2014b, 2011). Transformation in *S. pneumoniae* occurs following the quorum-dependent induction of “competence”, which is regulated by the secretion and detection of Competence Stimulating Peptide (CSP), encoded by *comC* (Håvarstein et al., 2006; Pestova et al., 1996). There are several alleles for *comC* and the gene encoding its cognate receptor, *comD*, although the vast majority of isolates carry either of two dominant mature signals encoded by *comC*, i.e. Csp-1 or Csp-2 (Evans and Rozen, 2013; Pozzi et al., 1996; Whatmore et al., 1999). Although these different allele combinations, referred to as pherotypes, are known to be mutually unresponsive, with isolates only responding to the CSP produced by isolates of their own pherotype (Iannelli et al., 2005), their role on the population structure of pneumococci remains unclear.

Two alternative hypotheses for the potential effects of multiple pherotypes on the population genetic structure of *S. pneumoniae* have been outlined. One hypothesis posits that because bacteria only bind and respond to their own CSP signal, pherotypes could ensure that bacteria only recombine with strains sharing the same pherotype, leading to a close association between pherotype and clonal structure (Håvarstein et al., 1997; Tortosa and Dubnau, 1999). In the second hypothesis, CSP-activated fratricide, whereby CSP-induced cells use secreted bacteriocins to kill uninduced cells, could alternatively facilitate recombination between strains with varying pherotypes. Importantly, this hypothesis assumes that strains of one pherotype preferentially kill strains of other pherotypes, after which the induced strains recombine with the DNA liberated from lysed cells (Claverys and Håvarstein, 2007; Claverys et al., 2006; Cornejo et al., 2010; Johnsborg et al., 2008). Although several recent studies have attempted to address the predictions of these models, results thus far are conflicting. While results from Carrolo et al. (Carrolo et al., 2014, 2009) support the idea that pherotypes facilitate within-pherotype recombination and therefore underlie population differentiation, results from Cornejo et al. (2010) are more consistent with the alternative. Importantly, the results of these previous studies were limited to moderately small samples of strains, and analyses were performed on a limited number of markers (partial sequences of seven MLST loci), thereby reducing the ability to infer genomic-scale patterns of recombination. To overcome these limitations, our aim here is to address this question using a bioinformatics approach based upon analysis of recombination rates within and between pherotypes from 4,089 full pneumococcal genomes.

## Materials and Methods

### Genomic data

We obtained 4,089 *S. pneumoniae* genomes from five publicly available sets (Supplemental Table 1): 288 genomes from GenBank, which include 121 genomes of pathogenic strains from Atlanta, Georgia, The United States (Chancey et al., 2015); 3,017 genomes of carriage strains from Myanmar refugees in Maela, Thailand (Chewapreecha et al., 2014); 616 genomes of carriage strains from Massachusetts, the United States (Croucher et al., 2013a); 142 genomes of carriage strains from Rotterdam, the Netherlands (Bogaert et al., 2001; Miller et al., 2016); and 26 PMEN (Pneumococcal Molecular Epidemiology Network) genomes (McGee et al., 2001; Miller et al., 2016). These genomes were previously assembled and underwent quality control (Miller et al., 2016). We located *comC* and *comD* within these genomes using an iterative DNA reciprocal BLAST search as previously described (Miller et al., 2016). We excluded Rotterdam strain 724 and Maela strain 6983_6#45 from all pherotype analyses because the genomes of both strains carry two complete *comC* alleles encoding for both Csp-1 and Csp-2.

### Phylogenetic analysis

To reconstruct the phylogenetic relationships of *comD*, we aligned full-length alleles using MUSCLE 3.8.425 (Edgar, 2004) and Geneious 7.1.5 (Kearse et al., 2012). After deleting sites with 5% or more gaps, we inferred the GTR+I+ γ substitution model using jModelTest 2.1.7 (Darriba et al., 2012). We used MrBayes 3.2 (Huelsenbeck and Ronquist, 2001; Ronquist and Huelsenbeck, 2003) for phylogenetic reconstruction across four independent runs.

The short length of *comC* prevented reconstructing the phylogenetic history with confidence; we instead used MUSCLE 3.8.425 (Edgar, 2004) and Geneious 7.1.5 (Kearse et al., 2012) to align amino acid variants and produce a UPGMA phylogram of amino acid identity.

In order to characterize the overall genetic population structure in *S. pneumoniae*, we used data from a previously reported full genome phylogenetic tree (Miller et al., 2016) based on alignment to strain R6_uid57859 (Hoskins et al., 2001). In summary, we used genome sites found in at least 99.5% of all the above *S. pneumoniae* genomes (1,444,122 sites) and used the single best maximum likelihood tree (ln(likelihood) of −3,2145,034.7) starting from fifteen unique random trees and from fifteen unique parsimonious trees, as calculated by RAxML v8.2.4 (Stamatakis, 2006) and ExaML v3.0 (Kozlov et al., 2015). This tree included 82 genomes from pneumococcus Complex 3 (Croucher et al. 2013) and 240 PMEN-1 genomes (Croucher et al., 2011); we excluded these strains from all of our analyses because they were originally sampled based on membership to clonal complexes and are therefore biased with respect to population structure and pherotype. 43 *Streptococcus sp.* viridans genomes were used as an outgroup for this tree, as previously reported (Miller et al., 2016).

For interspecific phylogenies, we identified *comC* and *comD* sequences from the available NCBI genomes for *S. infantis* (5 genomes), *S. mitis* (45 genomes), *S. oralis* (18 genomes), and *S. pseudopneumoniae* (40 genomes). The extensive sequence diversity in the mature CSP meant that we lacked confidence in aligning *comC*. Therefore, we used BAli-Phy v.2.3.6 (Suchard and Redelings, 2006) for phylogenetic analysis because it estimates both the alignment and the tree simultaneously. We used the HKY+*γ* model of substitution and constrained the leader sequence of *comC* to align together; BAli-Phy then estimated the full *comC* alignment and tree across eight independent runs using the S07 model of indel mutations.

Using MUSCLE 3.8.425 (Edgar, 2004) and Geneious 7.1.5 (Kearse et al., 2012), we aligned the full-length *comD* sequences and deleted sites with more than 5% gaps. For further analysis, we used the filtered polymorphic sites from Gubbins (Croucher et al., 2015); any regions in individual sequences with evidence of recombination from Gubbins were replaced with N’s. We used a GTR + *γ* model of nucleotide substitution as determined by jModelTest 2.1.7 (Darriba et al., 2012) to reconstruct the phylogeny using MrBayes 3.2 across four independent runs.

### Shannon Diversity

We located *comC* from 4 of 5 *S. infantis* genomes, 42 of 45 *S. mitis* genomes, all 18 *S. oralis* genomes, 39 of 40 *S. pseudopneumoniae* genomes, and 4,076 of 4,089 *S. pneumoniae* genomes. *comD* was also located in 4,040 *S. pneumoniae* genomes and from all other genomes except one *S. oralis* genome. We calculated Shannon diversity within each species based on amino acid identity across the entire gene. As each species had a different number of genomes, we calculated the Shannon diversity based on sub-sampling of 18 genomes for *comC* and 17 genomes for *comD*, corresponding to the minimum number of genomes in any species containing these genes, and we report the average of 1000 samples.

### Estimating population structure

We used hierBAPS (Cheng et al., 2013) on the full genome (2,038,615 bp) alignment to strain R6_uid57859 (Hoskins et al., 2001) in order to examine population structure. We used one run of 50, 60, 70, 80, 90, and 100 populations and four runs of 40 populations as upper bounds for the number of populations with between 2 and 4 levels of division. We found a constant number of 29-31 populations at the first level of division, and we used the aggregate of all runs at this level to divide genomes into populations for further analysis. We confirmed the inferred population structure using an orthogonal metric; for this, we measured nucleotide diversity across the entire genome by dividing the number of identical, non-gapped sites by the total number of non-gapped sites for pairwise combinations of genomes using Python 2.7.8. We then compared average nucleotide diversity between and within populations.

### Recombination events

We calculated the expected frequency of within- and between-pherotype recombination by assuming the null hypothesis that genetic exchange occurs randomly between pherotypes as a function of their respective frequencies. Accordingly, if i represents the frequency of strains producing Csp-1, the expected within Csp-1 recombination rate is *i^2^*. It then follows that the expected fraction of all recombination events that involve Csp-1 that occur strictly within-pherotype is

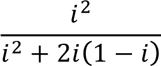

which reduces to:

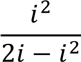

Similarly, the expected rate of between-pherotype recombination is *2ij*, where *i* and *j* represent the frequencies of the focal pherotypes.

We used GeneConv 1.81a (Sawyer, 1999) to detect recombination events of at least 100 bp between all pairwise comparisons of the 4,089 genomes; briefly, this program detects continuous sections of DNA in pairwise alignments that have higher identity than the surrounding regions after accounting for monomorphic sites. We analysed the genomes as ‘circular’, used Holm-Bonferroni correction for multiple testing (Holm, 1979), and used a Gscale constant of 2, which scales to the number of mismatches allowed while detecting recombination events.

We next calculated the fraction of observed recombination events relative to each pherotype. By dividing the observed recombination frequencies by the expected recombination frequencies and natural-log transforming this ratio, we derived an intuitive scaling: values greater than 0 indicate that there is more recombination than expected by chance, while values less than 0 indicate the opposite. We used the PropCIs package in R (R Core Team, 2013) to estimate 95% confidence intervals for this ratio, and we tested if the observed recombination frequency differed significantly from the expected recombination frequency using Pearson’s χ^2^ statistic using R (R Core Team, 2013). We additionally calculated observed and expected recombination frequencies by classifying strains by populations in place of pherotype.

To examine within and between pherotype recombination in more detail, we characterized the unique recombination events occurring between strains carrying the two dominant pherotypes (Csp-1 and Csp-2), which together comprise 95.5% of strains. By highlighting unique recombination events, the aim of this analysis was to remove the potentially biasing influence of vertical transmission, which
could cause more ancient recombination events to be overrepresented compared to newer events. In order to identify unique recombination events, we grouped events that shared an identical start and end position in the full genome alignment. Next, we calculated the average proportion of these unique recombination events taking place between strains with Csp-1 and Csp-2 for each group. We examined a range of cut-off points for the minimum proportion of strains that must be involved in each grouped recombination event, including: 0.05% of strains (≥2 strains, 1,663,312 events); 0.1% of strains (≥4 strains, 1,156,428 events); 0.5% of strains (≥20 strains, 381,084 events), 1.0% of strains (≥41 strains, 197,740 events), 2.5% of strains (≥102 strains, 59,980 events), and 5.0% of strains (≥204 strains, 19,428 events); this gradually focused the analysis on ancient recombination events. The expected proportion of between Csp-1 and Csp-2 recombination events was calculated as the expected encounter frequencies when only considering these two pherotypes, that is:

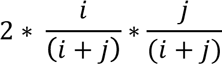

where *i* and *j* are the frequencies of Csp-1 and Csp-2, respectively.

## Results

### *comC* and *comD* diversity across *S. pneumoniae* genomes and geographic sites

We estimated pherotype frequencies from 4,089 *S. pneumoniae* genomes that include extensive sampling in four geographic sites (Table 1). The global population is dominated by two pherotypes (the mature signals encoded by the *comC* gene): Csp-1 (72.0% of strains) and Csp-2 (23.5% of strains). We also identified a novel variant in 2.1% of strains designated Csp-1_Short, which is identical to Csp-1 but with the final 4 residues deleted. Three other related variants contain between 1 and 3 “NFF” amino acid repeats, which we designate Csp-4_R1, Csp-4_R2, and Csp-4_R3, respectively. Note that Csp-4_R2 has previously been labelled Csp-4 while Csp-4_R3 is also called Csp-3 (Whatmore et al., 1999). 0.27% of strains contained no *comC* sequence, which may result from incomplete genome sequencing; the low fraction of genomes without a *comC* sequence indicates strong selection against phenotypes that do not produce CSP. The overall frequency ratio of Csp-1::Csp-2 in previous studies was approximately 73.7%::26.3% after minority pherotypes were removed (Carrolo et al 2009, Carrolo et al 2014, Cornejo et al 2010, Pozzi et al 1996, Vestrheim et al 2011); this is similar to the overall 75.4%::24.6% frequency ratio that we report after examining only Csp-1 and Csp-2 (Table 1). However, we found significant differences in Csp-1::Csp-2 frequency ratios between strains isolated from different geographic sites (Table 1), whose Csp-1 frequency ratios range from 77.6% to 65.9%. The Maela strains are significantly different from the Massachusetts and Rotterdam strains (p = 2.14 × 10^−9^ and p = 1.46 × 10^−8^ respectively, two-sample proportion test) but not significantly different from the Atlanta strains (p = 0.527, two-sample proportion test). The Atlanta strains are significantly different from the Massachusetts strains (p = 0.0351, two-sample proportion test), with both of these sets not significantly different from the Rotterdam strains (p = 0.193 and p = 1.00 respectively, two-sample proportion test).

**Table 1.**
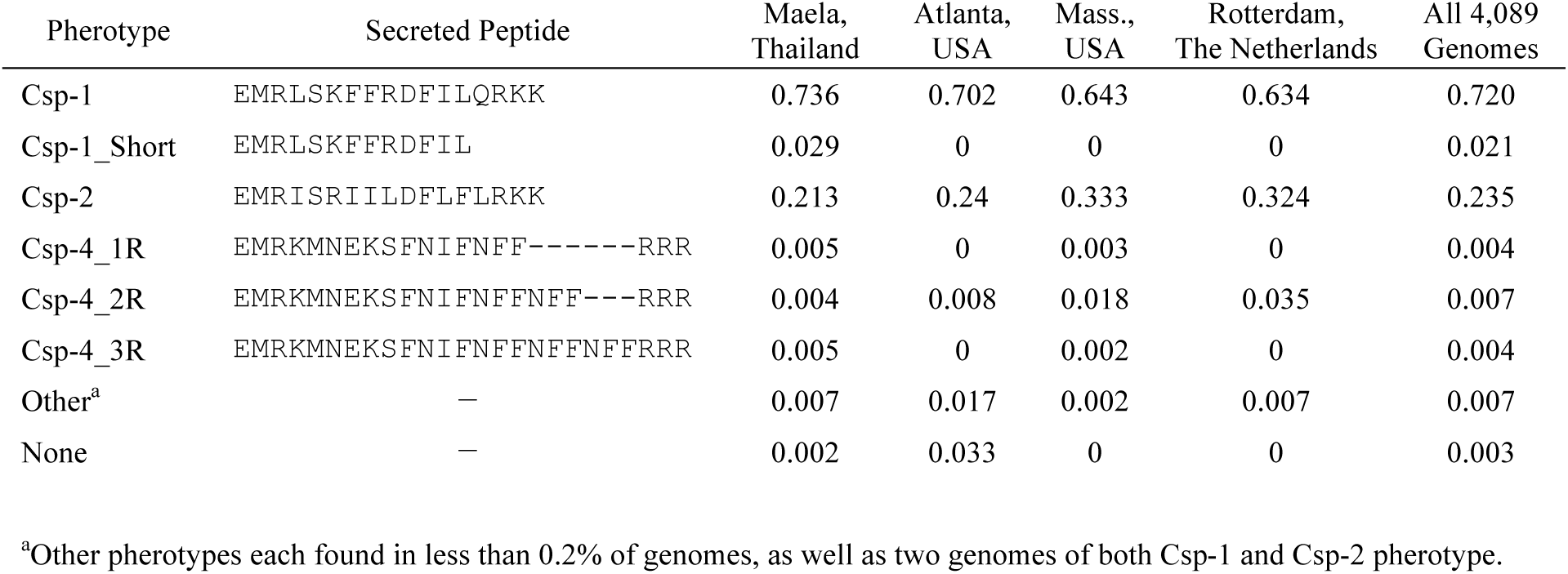
Frequencies of pherotypes within geographic sites.

The polymorphism in CSP is mirrored in the histidine kinase receptor for CSP, *comD*. We found excellent concordance between the pherotype each strain carries and its corresponding *comD* sequence (Figure 1). 935 of 936 strains with Csp-2 and full-length *comD* alleles contained *comD* alleles within a well-supported clade (posterior probability (PP) = 1.00) distinct from the other common (> 0.2% occurrence) pherotypes. Similarly, all *comD* alleles found with Csp-4 variants cluster in a well-supported (PP = 1.00) clade, with Csp-1 strains then forming a paraphyletic group with their *comD* alleles. The Csp-1 paraphyletic group also contains *comD* alleles associated with Csp-1_Short, which suggests this CSP may bind to the *comD* receptor similarly to Csp-1. This supports tight functional linkage within three groups of pherotypes and *comD* histidine kinases: Csp-1 (which includes Csp-1_Short), Csp-2, and the Csp-4 derivatives.

**Fig. 1.**
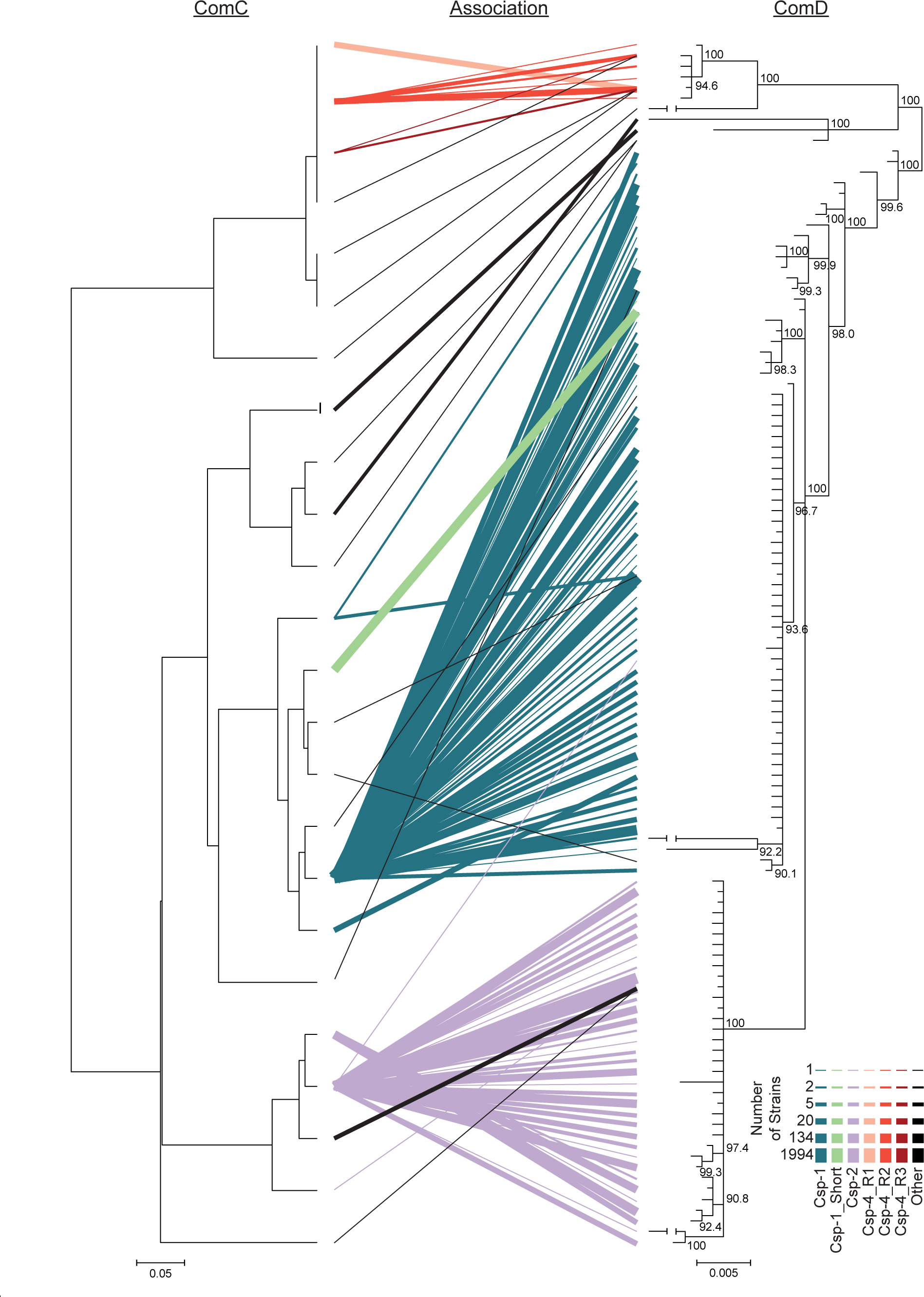
Associations between ComC and *comD*. The UPGMA clustering of ComC genes is shown next to the inferred phylogenetic relationship between *comD* alleles. ComC amino acid variants and *comD* nucleotide alleles found within the same genome are show as lines connecting the two phylograms, with thicker lines showing associations found in more strains. Line thickness is on a log scale. Classification of *comD* alleles is based on co-occurring CSPs within genomes, with a 99.8% correlation for the Csp-1 *comD* group, 99.2% correlation for the Csp-2 *comD* group, and 95.3% correlation for the Csp-4 *comD* group.

Interspecific gene trees of the signal gene *comC* (Supplementary Figure 1) lacked support for most interspecific clades. *comC* variants do not strictly cluster by species and Csp-4 derivatives form a clade with other non-pneumoniae *Streptococcus*, which could be indicative of: i) horizontal transfer, which is not uncommon among related *Streptococcus* species (Balsalobre et al., 2003; Duesberg et al., 2008; King et al., 2005); or ii) the possibility that trans-specific polymorphism is maintained in this locus as a balanced polymorphism (Gao et al., 2015; Ségurel et al., 2013). This second explanation is unlikely for *S. pneumoniae* given that 115 of 118 *S. pneumoniae* alleles form a well-supported clade that excludes other species in the interspecific *comD* phylogenetic tree of the histidine kinase receptor gene, in which we attempted to remove intragenic horizontal recombination (Supplementary Figure 2). By contrast, *S. mitis* and *S. pseudopneumoniae* freely intermix in all other clades, which is a pattern that supports trans-specific polymorphism between these two species. Shannon diversity of amino acid variants is lowest in *S. pneumoniae* compared to *S. mitis*, *S. oralis*, and *S. pseudopneumoniae* for comC (0.75 compared to 1.89-2.70), the mature CSP (0.67 compared to 1.17-2.63), and comD (1.51 compared to 2.66-2.78) (Supplementary Table 2). While diversity in *S. pneumoniae* is a product of the Csp-1:Csp-2 polymorphism, other viridians-group species appear to maintain higher diversity through a larger repertoire of CSP signal molecules (Supplementary Figure 1).

### Pherotype and population structure

We inferred the population structure of these strains using 10 independent hierBAPS runs with a full-genome alignment (Cheng et al. 2013; Figure 2); 23 of the resultant populations were invariant across all runs. The remaining genomes were assigned to one of 17 populations that each contained 21 genomes or more (0.5% of the total number of genomes) that co-occurred in all ten runs. 118 genomes (2.7%) could not be consistently classified into populations. Overall, this resulted in 40 estimated populations alongside the unclassified genomes. 25% of populations were not monophyletic on the full-genome phylogenetic tree, which is surprising but consistent with previous analysis of pneumococcal populations (Figure 2; Chewapreecha et al., 2014; Croucher et al., 2013b). Consistent with the hierBAPS analysis, we estimated less genome nucleotide diversity within a population than between populations for 39 out of 40 populations (mean of per-within population nucleotide diversity = 0.00404; mean of per-between population nucleotide diversity = 0.0111; p < 1.2 × 10^−57^ except for population 1, p = 0.059; corrected Wilcox test). Pherotypes are not equally distributed across populations (Figure 3A), with 20 populations fixed for a single pherotype. Similarly, geography has an unsurprising effect on population structure, as 16 of the 40 populations consist solely of strains collected in Maela, Thailand (Supplementary Table 3).

**Fig. 2.**
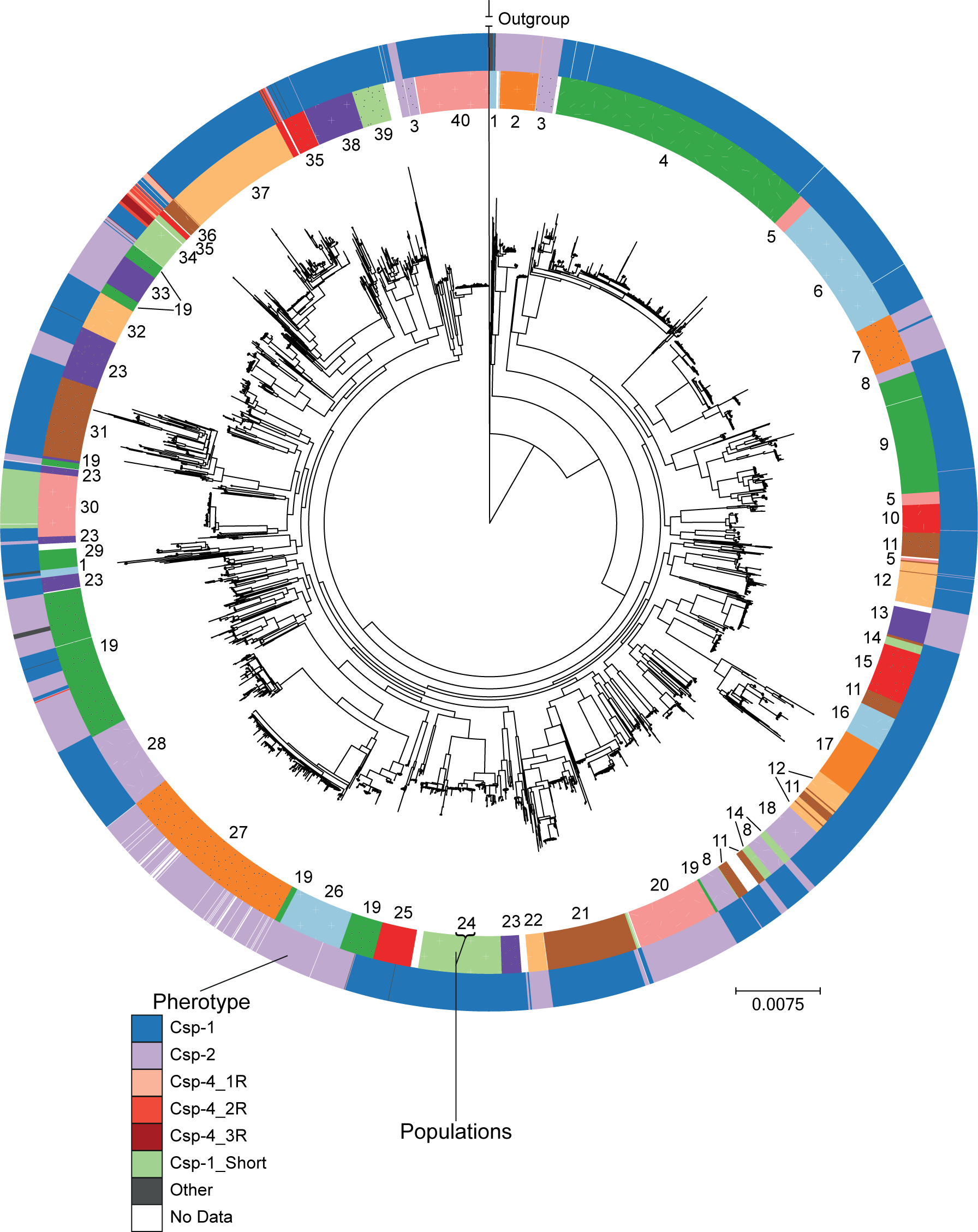
Phylogenetic relationship between *S. pneumoniae* genomes. The inner coloured ring shows the population grouping of strains as determined by hierBAPS, shown as a number and a colour/pattern. Genomes not classified into a population are white in the inner ring. The outer ring denotes pherotype.

### Recombination Events

To examine within versus between pherotype recombination frequencies, we compared the observed fraction of recombination events to the expected fraction of recombination events that assumes strains recombine randomly. Overall, we found no evidence that recombination is limited between populations. While populations 9 and 21 had significantly more within-population recombination events than expected (Figure 3B), 37 other populations had significantly less within-population recombination events than expected (p ≤ 1.09 × 10^−6^ except for population 1, p = 0.395; Newcombe proportion test with Holm-Bonferroni correction). The pherotypic diversity of populations, as estimated by Shannon diversity, created an interesting pattern with the proportion of observed between-population recombination events (Figure 3C), although a linear relationship is not statistically significant (Spearman’s rank correlation ρ = 0.254, p = 0.280 after removing populations with Shannon diversity = 0).

**Fig. 3.**
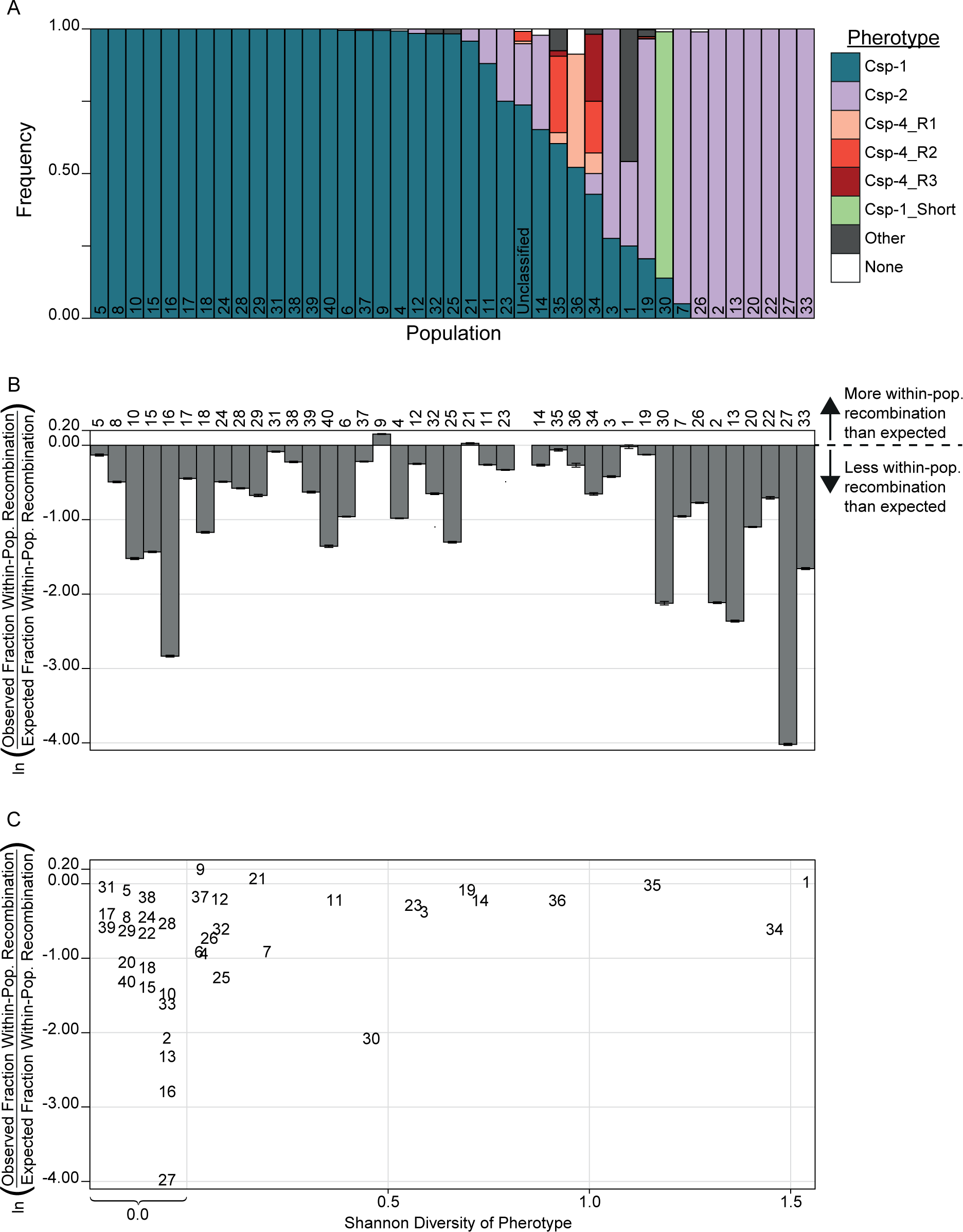
Relationship between pherotypes and populations. A) Distribution of pherotypes within each population. B) Observed / expected fraction of within-population recombination for each population, with values below zero indicating less observed within-population recombination than expected. Error bars show 95% confidence intervals. C) The same recombination ratio as part B with populations’ Shannon diversity of pherotypes, with populations shown as numbers.

Figure 4 shows estimates of recombination for all common (> 0.2% of strains) pherotype combinations, with all observed recombination frequencies differing significantly from those expected (p < 4.9 × 10^−6^; Newcombe proportion test with Holm-Bonferroni correction; Figure 4A). Of the 30 between-pherotype comparisons, 15 estimates of recombination frequencies were significantly higher than expected, and 15 estimates were observed significant less frequently than expected. Importantly, we found a higher observed recombination frequency than expected between strains expressing the two dominant pherotypes Csp-1 and Csp-2 (p < 10^−99^; Newcombe proportion test with Holm-Bonferroni correction). In total, recombination between these two pherotypes comprises 34.7% of all recorded recombination events; thus an excess of recombination for these pairs implies that the general role of pherotypes is to marginally increase between-pherotype recombination rates. Five of the six within-pherotype observed recombination frequencies are lower than expected, with Csp-4_R2 as the only exception (Figure 4A); three of these frequencies are the lowest of all pherotype combinations within their respective pherotype.

**Fig. 4.**
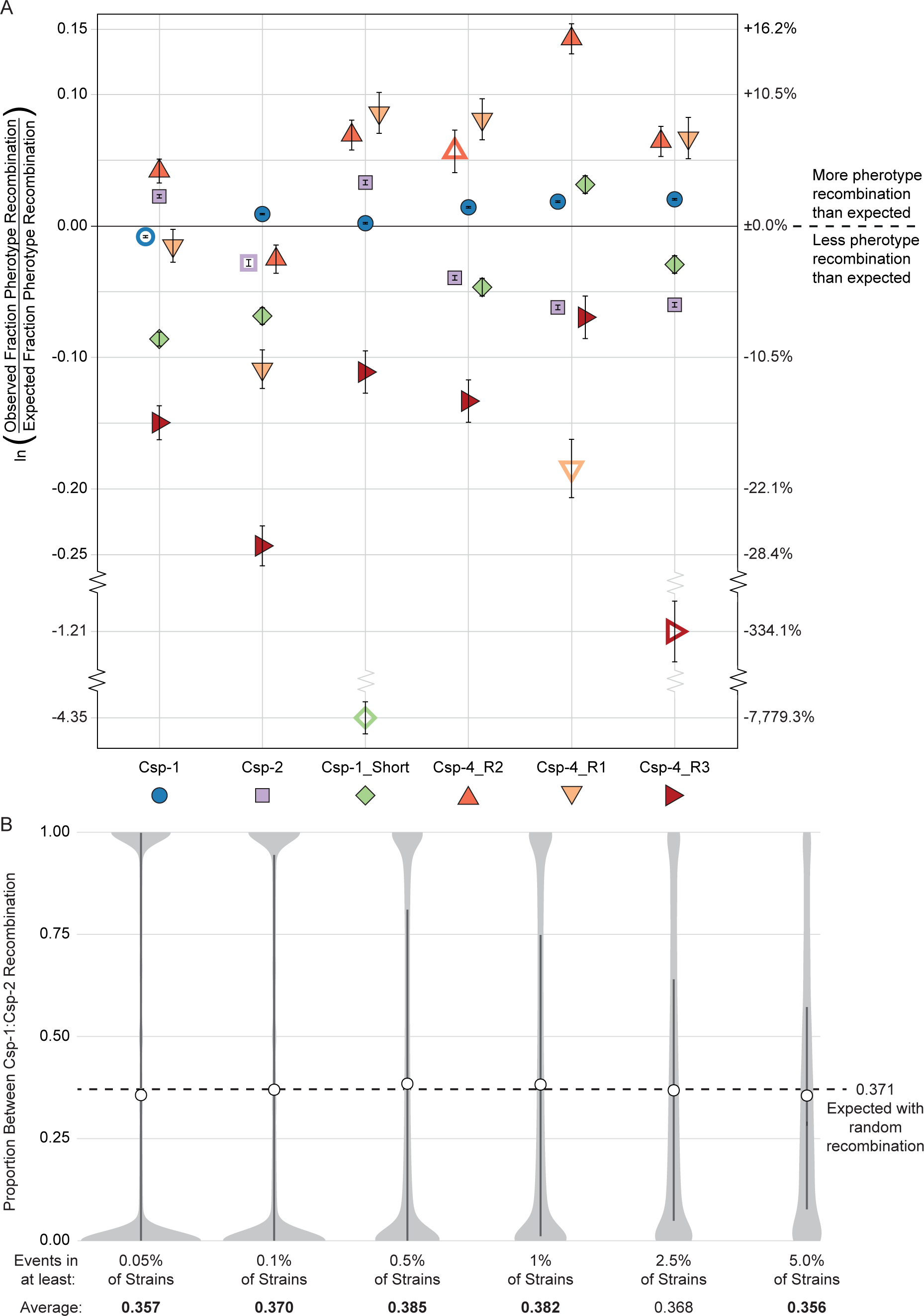
Recombination between and within pherotypes. A) Observed / expected fraction of recombination within and between pherotypes. Empty shapes are within-pherotype comparisons, while filled shapes show between-pherotype comparisons. Colours correspond to pherotypes as in Figure 1. Error bars show 95% confidence intervals. B) Distribution of the proportion of recombination events between Csp-1 and Csp-2, in which all recombination events with identical breaks points in the full-genome alignment are grouped as a single event. Light grey shapes show density estimate; dark bars incorporate the 25^th^ to 75^th^ percentile; white circles indicate the average proportion. Averages in bold are significantly different from the null expectation.

A potential caveat of these results is that we consider every pairwise recombination event as independent. However, recombination events could have occurred at any time during the ancestry of these strains and then persisted via vertical descent, which may lead to biased estimates of recombination frequencies if a single, historic recombination event is counted in each of the multiple strains that descended from their common ancestor. As the true number of recombination events through history is unknown, we considered the opposite extreme, in which pairwise recombination events with identical start and end positions in the full-genome alignment originated from a single recombination event. We focused only on Csp-1 and Csp-2 pherotypes, and we considered unique recombination events found in a range of at least 0.05% of genomes (≥2 genomes) to at least 5.0% of genomes (≥204 genomes) in order to focus on more ancient recombination events (Figure 4B). The mean proportion of between Csp-1::Csp-2 events of five out of six distributions are significantly different than a null expectation based on pherotype frequencies in the global population (p < 0.0253; Wilcox test for difference to μ = 0.371); however, no values differ by more than 4.0% from the null expectation of 0.371, with three averages significantly above and two averages significantly below the null value. This result indicates that any potential bias introduced by considering all recombination events independently is, at most, marginal.

## Discussion

Several studies, including this one, have found that the two dominant pherotypes of the quorum-dependent regulator of competence in *S. pneumoniae* (i.e. Csp-1 and Csp-2) are maintained at relative frequencies of roughly 70:30 (Carrolo et al 2014; Vestrheim et al 2011; Cornejo et al 2010; Carrolo et al 2009; Pozzi et al 1996). Our results also indicate that there is subtle geographic variation in the ratio of these dominant pherotypes, ranging from 66:34 to 78:22 (Table 1). These results naturally lead to questions of how this pherotype ratio is maintained (to the near exclusion of other pherotypes), especially at similar, yet not identical, levels in disparate geographic sites. The null explanation is that pherotypes are neutral with respect to bacterial fitness, although three lines of evidence counter this explanation. First, other studies suggest that *comC* is evolving under positive selection (Cornejo et al 2010; Carrolo et al 2009). Second, other *Streptococcus* viridans species have a higher diversity in CSP, ComC, and ComD; this diversity is caused by a large number of pherotypes within each species as opposed to the two dominant pherotypes in *S. pneumoniae*. Third, each geographic site is comprised of strains from anywhere between 14 and 36 different populations (Supplemental Table 3), yet each site maintains the approximate 70:30 ratio of major pherotypes. This inter-population pattern between geographic sites is unlikely to occur through neutral mechanisms.

Irrespective of the factors maintaining pherotype polymorphism, what are the consequences of their maintenance for pneumococcal populations? Two divergent hypotheses have been explored to answer this question. One holds that these variants reinforce population subdivision by restricting recombination to strains that carry the same pherotype (Carrolo et al 2009). The alternative hypothesis suggests that lysis of non-identical pherotypes by the process of fratricide leads to inter-pherotype transformation and therefore the elimination of pherotype-specific population structuring (Cornejo et al 2010). Our data, based on the analysis of recombination from more than 4,000 full genome sequences, are not clearly consistent with the extreme version of either hypothesis. We find a slight tendency towards increased between-pherotype recombination for the dominant pherotype classes (Csp-1 and Csp-2; Figure 4A), and these results clarify that pherotypes are not a barrier to recombination in this species. However, these data are not without limits, in particular the difficulty of inferring recombination among closely related strains, which could partly explain the lower frequencies of within-pherotype (Figure 4A) or within-population (Figure 3B) recombination. Because this should only have minimal influence on estimates of between-pherotype recombination frequencies, as strains carrying different pherotypes tend to come from different populations (Figure 3B), we do not view this is a bias that is likely to influence our conclusions.

*S. pneumoniae* live in surface-associated biofilm communities, where in the case of single-strain colonization, they will be surrounded by clone-mates. If some of these cells become competent and lyse the others via competence-induced fratricide (i.e. induced production of bacteriocins that target uninduced cells), there will be little signature of this event at the genomic level. However, it is increasingly clear that multiple-strain infections are more common than previously thought; co-colonization within the nasopharynx is observed in up to 50% of individuals (Wyllie et al 2014; Rodrigues et al 2013), and because rare variants are less likely to be detected during sampling, this may be an underestimate of the true occurrence of co-colonisation. In these cases, there is opportunity for genetic exchange between different genotypes. If the co-infecting genotypes express the same pherotype, both will respond to the same peptide signal and both genotypes could, in principle, release DNA that will be available to the other. However, if the genotypes express different pherotypes, then the genotype that first initiates competence may lyse the other non-responding genotype via competence-induced fratricide, leading to more unidirectional uptake. While recombination during single pherotype infections could reinforce any pre-existing association between pherotype and population structure, recombination during mixed-pherotype infections would cause this association to decline or disappear. Our results are most consistent with this latter scenario.

How often do mixed pherotype colonization events occur? If the likelihood of colonization is random with respect to pherotype, then the probability that both pherotypes will be found to co-occur is simply twice the product of these relative frequencies. In two recent studies where this has been measured, mixed pherotype infections are common in cases where multiple strains are seen (47.5% and 57.1% in strains from Portugal and Norway, respectively) and do not vary from the null expectation of random colonization (Valente et al 2012, Vestrheim et al 2011). These results have several important implications. First, they suggest that fratricide during the induction to competence has minimal, if any, impact on the within-host competitive dynamics of co-occurring strains. Second, they suggest that opportunities for recombination between strains expressing different pherotypes are widespread. Accordingly, we would predict pherotypes to have little direct influence on population structure, a prediction borne out by the results presented here and elsewhere (Figure 3A; Cornejo et al 2010). This, of course, does not preclude structure arising from other influences, e.g. serotype specific immunity, differences in antibiotic resistance or other attributes leading to biases in colonization. However, pherotypes neither appear to underlie this structure nor to eliminate it due to ubiquitous inter-pherotype recombination (Figure 4A).

A final explanation for a fixed pherotype ratio is that diversity is maintained for reasons wholly distinct from their effects on competence. In addition to competence, CSP induces more than 150 pneumococcal genes, and only a small fraction of these are required for DNA uptake and recombination (Peterson et al 2004; Peterson et al 2000). Different concentrations of CSP are required for competence activation between Csp-1 and Csp-2 with their respective receptors (Carrolo et al., 2014; Iannelli et al., 2005), which determine the population density at which CSP induces these genes and could drive large phenotypic differences between pherotype based only on *comC* and *comD* variation. We were unsuccessful in finding genomic loci that co-associates with CSP via selection using a genome-wide association study with pherotype as a phenotype, which suggests that any ecological differentiation between phenotype may be caused only by differences in *comC* and *comD*. However, Carrolo et al. (2014) find striking differences in the ability for strains expressing Csp-1 and Csp-2 to form biofilms, with Csp-1 strains producing biofilms with greater biomass. Although the authors suggest that this difference may lead to differences in the colonization success and transmissibility of isolates, epidemiological data discussed above are not consistent with this possibility; co-colonization with multiple pherotypes occurs as often as expected by chance (Valente et al 2012, Vestrheim et al 2011). An alternative, which could lead to a form of balancing or frequency-dependent selection maintaining pherotypes at intermediate frequencies, is the possibility that differences in biofilm-associated biomass that influence stable colonization may trade-off with reductions in transmissibility. By this scenario, Csp-1-expressing strains may form more robust biofilms during colonization, while strains expressing Csp-2 are more proficient at dispersal. Although testing these possibilities is beyond the scope of this work, these ideas can potentially be examined empirically in both *in vivo* or *in vitro* models. Furthermore, they make the prediction that carriage duration should vary as a function of pherotype, a possibility that could potentially be retrospectively examined from epidemiological studies.

Although our results are unable to clarify the evolutionary factors that lead to the origin and maintenance of pherotypes in *S. pneumoniae*, our comprehensive analysis demonstrates the long-term effects of this polymorphism on recombination in this species. In summary, while pherotypes can apparently facilitate recombination between the major pherotype classes, this effect is weak and has no evident impact on population structure of this pathogen. Explanations for pneumococcal population structure therefore lie outside of pherotypes, and indeed, explanations for pherotypes may lie outside of their effects on recombination.

## Acknowledgements

Bioinformatic work was carried out on the Dutch national e-infrastructure with the support of SURF Foundation. This work made use of the facilities of N8 HPC Centre of Excellence, provided and funded by the N8 consortium and the Engineering and Physical Sciences Research Council (grant number EP/K000225/1). The Centre is co-ordinated by the Universities of Leeds and Manchester. This work was supported by the Biotechnology and Biological Sciences Research Council (grant number BB/J006009/1) to DER and ISR and by the Wellcome Trust (105610/Z/14/Z) to the University of Manchester.

